# Identifying regulators of aged fibroblast activation in 3D tissue models

**DOI:** 10.1101/2025.02.10.636275

**Authors:** Hui Liu, Luezhen Yuan, G.V. Shivashankar

**Author notes:** Author contributions H.L. and G.V.S. designed the project. L.Y. analyzed Prize Collecting Steiner Tree. H.L conducted and analyzed other data. H.L., L.Y. and G.V.S. wrote the manuscript. All authors have read, edited and approved the manuscript. Competing Interest Statement: All authors have no competing interest.

## Abstract

Robust 3D tissue culture models to activate aged fibroblasts for cell-based therapies and identify regulators of such activation are still missing. In our previous study, we showed that aged fibroblasts can be activated simply through applying compressive force, without the need for exogenous factors, leading to increased migration. In this study, we develop a pipeline to evaluate the role of specific pharmacological inhibitors for transcription factor inference and cell migration involved in aged fibroblast activation. By integrating RNA-seq data with bioinformatic tools (prize collecting Steiner tree method and iRegulon) we inferred 15 candidates. In addition, we used cell migration and heterochromatin content as readouts for validating these candidates. Furthermore, we identified three potential master regulators of fibroblast activation and rejuvenation: FOXO1, STAT3, and PDK1. These findings offer valuable insights for future drug discovery, disease modeling, and regenerative medicine.

## Introduction

Cellular states can be influenced by their local microenvironment, including biochemical signals and physical cues (1–3). Several studies have demonstrated that physical forces can enhance reprogramming efficiency or even induce somatic cells to adopt a stem-like state without the need for exogenous factors (4–6). It is well-established that cells exhibit plasticity and can modify their shape and behavior in response to external microenvironmental cues through the mechanosensor–mechanotransduction system, leading to changes in gene expression (7, 8). In our previous study, we showed that aged human fibroblasts can be activated or rejuvenated under compressive forces (71). However, the underlying mechanisms behind this phenomenon remain unknown. Therefore, in this study, we aim to explore the molecular mechanisms underlying mechanical stimuli induced transcriptional regulation of aged fibroblasts activation.

Transcription factors (TFs) play a crucial role in controlling cell identity and function (9). Identifying the key TFs involved in mechanical force-induced rejuvenation is important for understanding this process and for developing potential therapeutic targets which may benefit healthy aging. TFs are proteins that can be located in the cell membrane, cytosol, or nucleus (10). These proteins bind to specific DNA sequences via promoter or enhancer regions to initiate the transcription process (11). They regulate gene expression through diverse mechanisms, such as integrating signals from signaling pathways, interacting with RNA polymerase, recruiting co-activator or co-repressor proteins, and acting as bridges for chromatin remodelers (12). Notably, some TFs are mechanosensitive, such as YAP/TAZ, P53, MRTF, P65 and STAT3 (13–18). This gives us hints that certain TFs may play a key role in our previously established model under compressive force conditions, potentially acting as mediators of the cellular response and contributing to the rejuvenation process.

Based on our established 3D cell culture model and RNA-seq data, we applied the Prize-Collecting Steiner Tree and iRegulon methods to identify fifteen key master TF regulators. Following this, we performed selective TF inhibitor screening and used cell migration and heterochromatin content as readouts. FOXO1, STAT3, and PDK1 were identified using this assay. This pipeline and findings offer valuable insights into the fields of anti-aging research, drug discovery, and regenerative medicine.

## Materials and Methods

### Cell culture

GM08401 (75 years old) healthy human dermal fibroblast cells (HDFs) (male origin) were obtained from the NIGMS Human Genetic Cell Repository at the Coriell Institute for Medical Research. The HDFs were cultured in MEM (Gibco, 11090-081) with 15% FBS (Thermo Fisher, 16141079), 1% P/S (Penicillin and Streptomycin) (PAN BIOTECH, P06-07300), 1% Glutamax (100x, Gibco, 35050-038) and 1% NEAA (100x, Gibco, 11140-035) under 5% CO2 and 37 °C. Formation of spheroid and application of PDMS and static compressive force, using 7 stacked coverslips, in 3D spheroid model is the same as our previous work (71).

### Drug screening

All drugs, along with their storage conditions and final concentrations, are listed in Supplementary Table 1. After spheroids were formed overnight on fibronectin-coated patterns, the medium was carefully removed, and 1 mL of fresh medium containing inhibitors was added. Following a 1-hour incubation, 400 μL of collagen type I containing the complete medium without inhibitors (working concentration 1 mg/mL, Gibco, A1048301) was carefully layered on top of the spheroids. Once the collagen solidified after 1 hour incubation, a glass ring, 7 stacked coverslips (to apply compressive force), and 2 mL of medium with inhibitors were added. The samples were then cultured for an additional two days. Finally, the collagen gels were processed for immunostaining and imaging.

### Image acquisition and analysis

At the two-day time point, samples were fixed with 4% PFA (Merck, F8775-25ML) for 1 hour. Coverslips were carefully removed using bent needle tips and tweezers. A 400 μL mixture of PBS containing DAPI (Thermo Fisher Scientific, R37605, 1 drop per 1 mL) and ActinGreen (Thermo Fisher Scientific, R37110, 1 drop per 1 mL) was added to each Ibidi dish and incubated overnight at 4°C, protected from light. All images were captured using the EVOS M5000 (Thermo Fisher Scientific, 4x objective) and the Nikon Ti2 Confocal Imaging System (20x objective). Heterochromatin analysis (i80_i20) code (images came from confocal images) comes from https://github.com/GVS-Lab/chrometrics.git.

### Prize Collecting Steiner Tree analysis

RNAseq data came from our previous work (71). Network analysis was done similarly as described previously (19). We adapted this network optimization method to model ERK dependent differential expressed genes responding to load (20). One input of this method is an interaction network with assigned values on nodes and edges. To construct this interaction network, human protein-protein interaction data were sourced from the STRING database (version 11.5), encompassing 17,804 genes and 937,906 interactions (21). Relationships between transcriptional regulators and target genes were obtained from the hTFtarget database and various online datasets (downloaded from Harmonizome and Enrichr database as described in previous publication) (19, 22–25). This compilation included 774 transcriptional regulators, 18,932 protein-coding genes as targets, and 4,419,504 interactions.

Transcriptional regulators with protein-protein interaction information were designated as transcriptional regulator nodes (referred to as TF nodes), while other proteins in the protein-protein interaction data were categorized as upstream protein nodes (referred to as Protein nodes). Target genes of the transcriptional regulators were denoted as RNA nodes. The costs on the edges of the network, representing interactions between nodes, were calculated as -log(score/1000), with the score derived from the quality of links between proteins stored in the STRING database (multiplied by a factor of 1000 upon download). Edges linking transcriptional regulators and targets were assigned a cost of zero.

The values (or Prizes) of target (RNA nodes) were determined by the averaged absolute log2 fold change of each common gene (199 in total) 1) upregulated in the load group (7 stacked coverslips) compared to the control (C) group and 2) downregulated in the PD98059 treated group compared to the static compression group. These genes were likely upregulated under the load condition and influenced by ERK signaling, as evidenced by their downregulation under PD98059 treated conditions. The log2 fold change of genes upregulated in the load group compared to the C group (a total of 278 genes) was assigned to the values of corresponding Protein or TF nodes. Network optimization, utilizing the Prize-Collecting Steiner Tree method, was then executed to derive a regulatory network connecting potential upstream protein and transcriptional regulator nodes with their targets. ERK-dependent transcription factors were identified based on their expression changes under compression and the number of targets they can regulate.

The iRegulon plug-in in Cytoscape software was also used to identify additional potential TFs (26). TFs with a normalized enrichment score (NES ≥ 3.16).

### Statistical analysis

All plots and statistical analysis were performed with Origin 2024. Unpaired, two-tailed student-t test was used to compare two groups. For box-and-whisker plots: The box represents the interquartile range (IQR), encompassing the middle 50% of the data. The bottom of the box marks the first quartile (25th percentile), and the top marks the third quartile (75th percentile). The line inside the box indicates the median (50th percentile). The whiskers extend to the smallest and largest values within 1.5 times the IQR, while outliers are represented by asterisks. All illustration figures are generated in Biorender.

## Results

### Pipeline for selective TF screening and migration behavior as a phenotypic readout

In previous work, we established a 3D cell culture model under static load to examine whether fibroblast activation can be measured by their migration behavior. Such cell activation was demonstrated using two loading conditions: 1× (three stacked coverslips) and 2× (seven stacked coverslips). The results showed that the 2× load generated a higher activation response. In addition, we identified the ERK dependent pathway as one of the major regulators of force induced activation, using the ERK inhibitor PD98059. To identify potential regulatory mechanisms underlying this process, we developed the following pipeline as in Figure 1A. We first identified a list of 119 differentially expressed genes (DEGs) (Figure 1B) that were upregulated in the static compression group compared to both the unload group and the PD98059 group. This analysis provided a list of ERK-related genes, and therefore, we developed a strategy to identify potential transcription factor (TF) targets to perform a small scale drug screening. Cell migration level as a readout provides a quick method to identify critical intermediates by using DAPI and actin staining, bypassing the need for time-consuming antibody staining. In addition, the DAPI images provide a measure of the chromatin states, as established in our previous work (72). Since heterochromatin appears brighter than euchromatin across the nucleus, our pipeline also links the inhibitor effects both on cell migration behavior and chromatin organization. Ultimately, this approach allowed us to identify the potential upstream regulators controlling the fibroblast activation.

**Figure 1.**
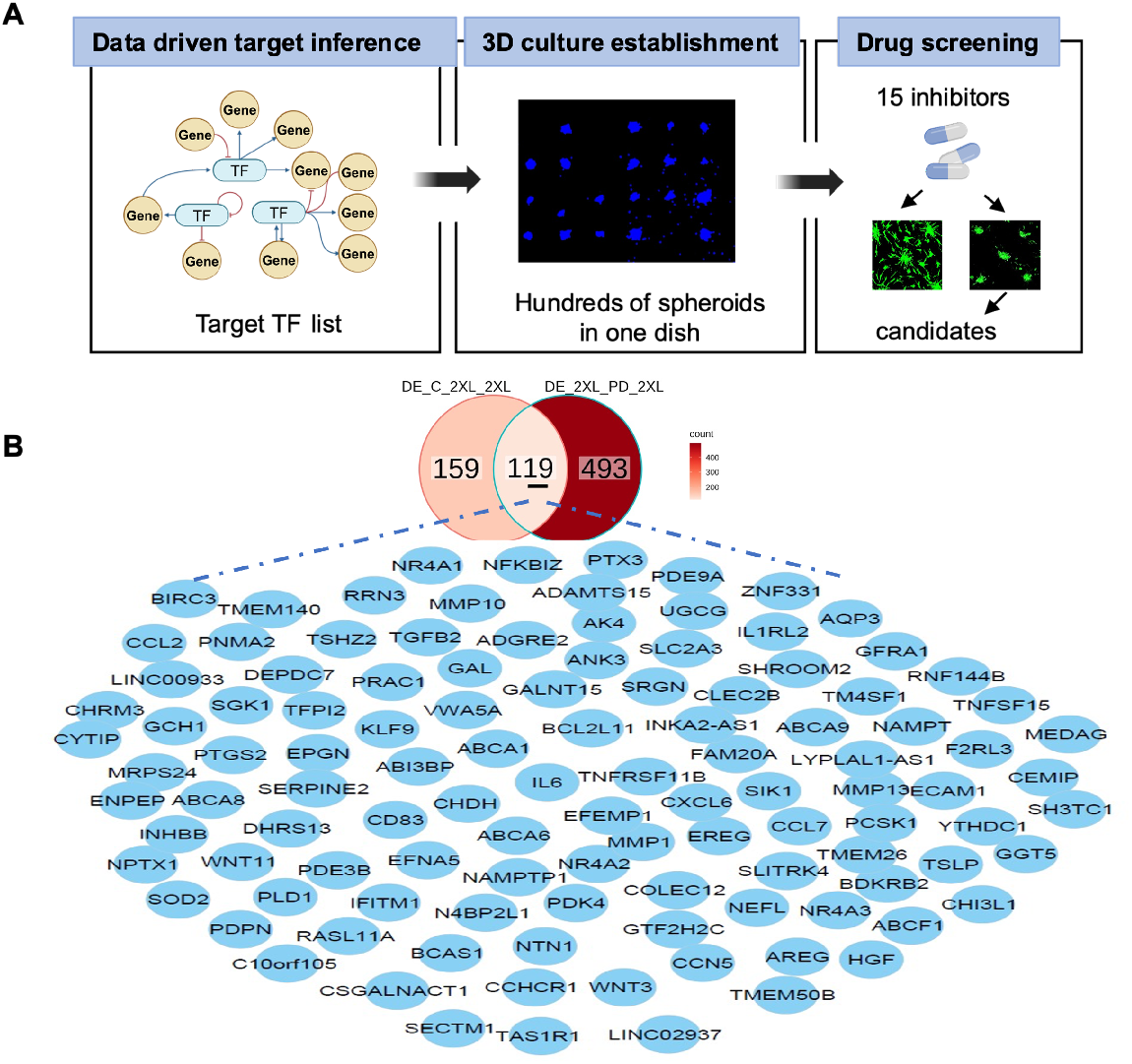
Schematic of selective inhibitors screening. (A) Pipeline for drug screening, including data driven transcription factor (TF) inference from RNAseq data, 3D spheroid culture as drug screening platform, cell migration as a main readout. (B) 119 DEG list from RNAseq data comparing control with load group, load group with ERK inhibitor (PD98059 treated group). DE: differential expression; C: unload group; 2XL: 2xload group; PD: ERK inhibitor PD98059 treated group.

### Construction of the Protein-TF-Target gene Regulatory Network by Prize-Collecting Steiner Tree method

To identify potential regulatory mediators of the ERK-mediated signaling pathway under compressive force conditions, which could act as candidates of targets for above mentioned drug screening pipeline, we applied the Prize-Collecting Steiner Tree analysis to construct the Protein-TF-Target gene network (Figure 2A). In this method, potential transcriptional regulatory pathways were modeled using genes both upregulated under load and downregulated under ERK inhibitor PD98059 treatment (119 DE gene list). The backbone interaction networks were sourced from the STRING PPI database, transcription factors (TFs) were obtained from the hTF target database (as described in detail in method). Based on the gene expression levels of the identified transcription factors (TFs) in the network we found that the top six differential expressed TFs were NR4A1, CEBPD, KLF9, ZNF331, IRF1, and NR6A1 (Figure S2A). Several TFs in the inferred transcriptional regulatory network have a high number of targets within the 119 gene list, as shown in Figure 2B. All TFs identified in the table are potentially involved in regulating the genes within the 119 DE gene list.

**Figure 2.**
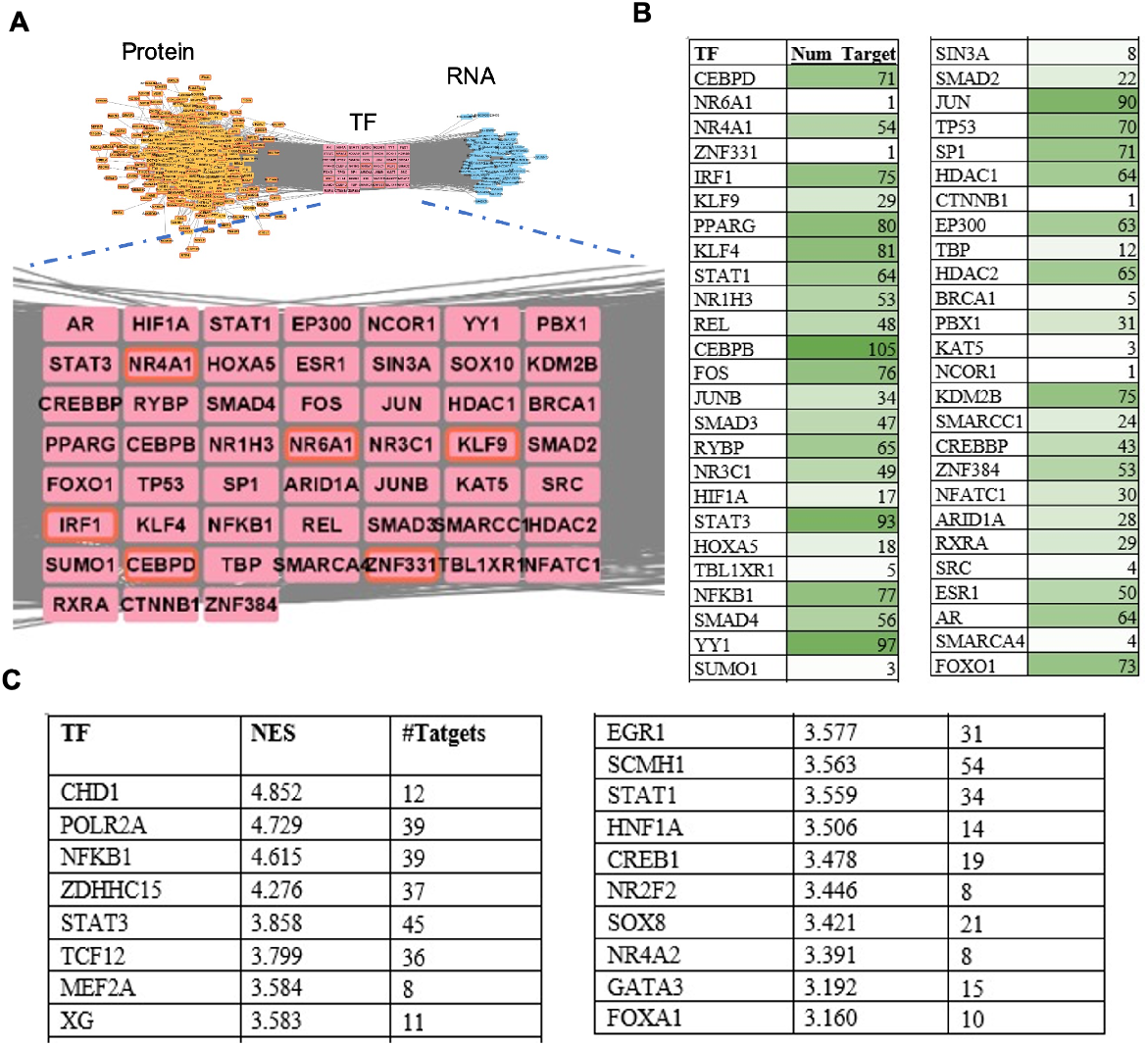
Inferring key transcription factors from differential expressed gene list. (A) Transcriptional regulatory network derived from Prize-Collecting Steiner Tree method, optimized for selected differential expressed genes (DEG). Yellow nodes represent upstream of TFs (Protein node); Pink nodes represent TF; Blue nodes represent 119 DEG (RNA node). (B) Table for numbers of potential targets for all TFs found by Prize-Collecting Steiner Tree method. Dark green color means the number of target genes is high. (C) Table for all enriched TFs found by iRegulon method.

We also employed iRegulon as an additional method to predict TFs. iRegulon is a tool designed to infer enriched TFs when RNA-seq data is available, but ChIP-Seq data is not (26). It integrates over nine thousand position weight matrices (PWMs) and operates through a rank-based motif discovery and motif-to-TF mapping procedure. After inputting 119 DEG, we got a list of potential TFs. Figure 2C presents a table listing all TFs identified by the iRegulon plugin, including their Normalized Enrichment Score (NES) and the number of target genes. Therein, Figure S2B illustrates the regulatory network between representative transcription factors STAT1, STAT3, GATA3, MEF2A, CHD1, and their target genes. Notably, all these TFs, except STAT1, target NR4A1, which was identified in our Steiner tree analysis and has been shown to be associated with rejuvenation (28).

### Selective transcription factor inhibitor screening and pathway analysis

After compiling the list of transcription factors (TFs), we selected 15 inhibitors for the current analysis. Based on their primary targets, we categorized them into four groups: NR4A1, STAT3, GATA3, NFATC1, FOXO1, TP53, STAT1, and JUN belong to transcription factors; SRC, SMAD3, and PDK1 are classified as signal transduction molecules; SIN3A, HDAC2, and NCOR1 are chromatin modifiers and co-regulators; and HIF1A is grouped under hypoxia and stress response.

According to the signaling pathways they are involved in, TP53, JUN, and STAT1 are associated with the NF-κB pathway; STAT1 and STAT3 with the JAK-STAT pathway; SMAD3 with the TGF-β/SMAD pathway; HIF1A with the hypoxia pathway; NR4A1 with nuclear receptor signaling; NFATC1 with the calcium signaling pathway; PDK1 and FOXO1 with the PI3K/Akt pathway; JUN with the MAPK/ERK pathway; SRC with the Src family kinases pathway; and TP53 with the p53 pathway.

Using the 3D cell culture model described in our previous work (see method section), we performed a cell migration assay with the 15 inhibitors to assess aged cell activation. As shown in Figure 3A (large area as shown in Figure S1), cell migration behavior varied across these inhibitor groups. Figure 3B highlights that STAT3, FOXO1, and PDK1 inhibitor groups significantly inhibit cell migration. We then examined their pathway involved (Figure 4C) and found that PDK1, FOXO1, and STAT3 are downstream of SRC. PDK1 and FOXO1 are involved in the PI3K/AKT signaling pathway, while STAT3 is involved in the JAK/STAT pathway. Notably, our findings have narrowed the key regulators to two signaling pathways: PI3K/AKT and JAK/STAT.

**Figure 3.**
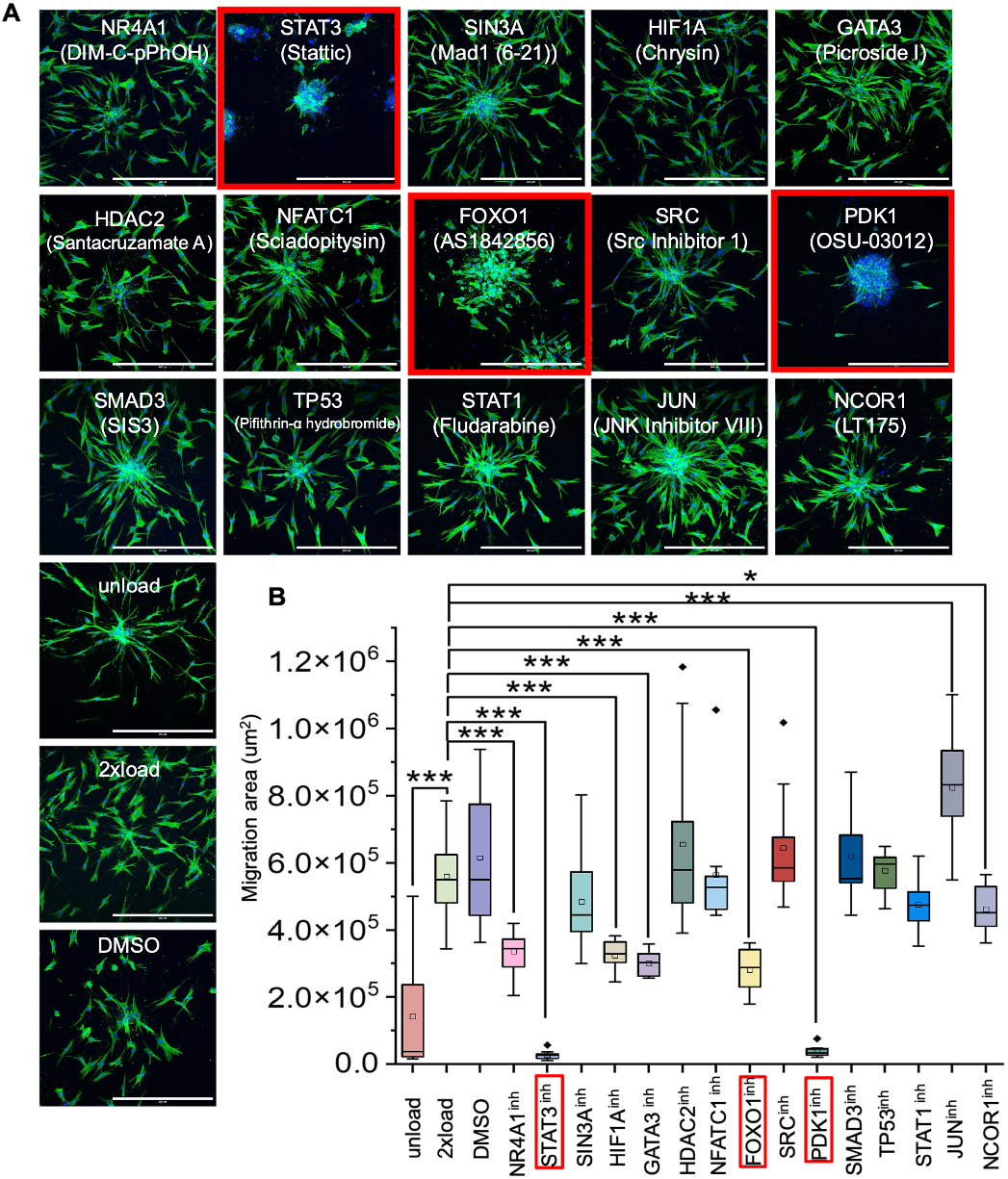
Selective TF inhibitor screening using cell migration as a readout. (A) Representative immunofluorescence confocal images to check spheroid spreading area. Nucleus is labeled in blue. F-actin is labeled in green. (Scale bar, 500 μm). (B) Quantification data of the spread area of the spheroid (n=10 spheroids per condition from EVOS data). All the experiments were repeated at least three times independently with similar results. P values in Figure (B) were calculated by unpaired, two-tailed Student’s t test. Other groups are compared to the 2XL group. *P<0.05; **<0.01; ***P<0.001; No asterisks: not significant.

**Figure 4.**
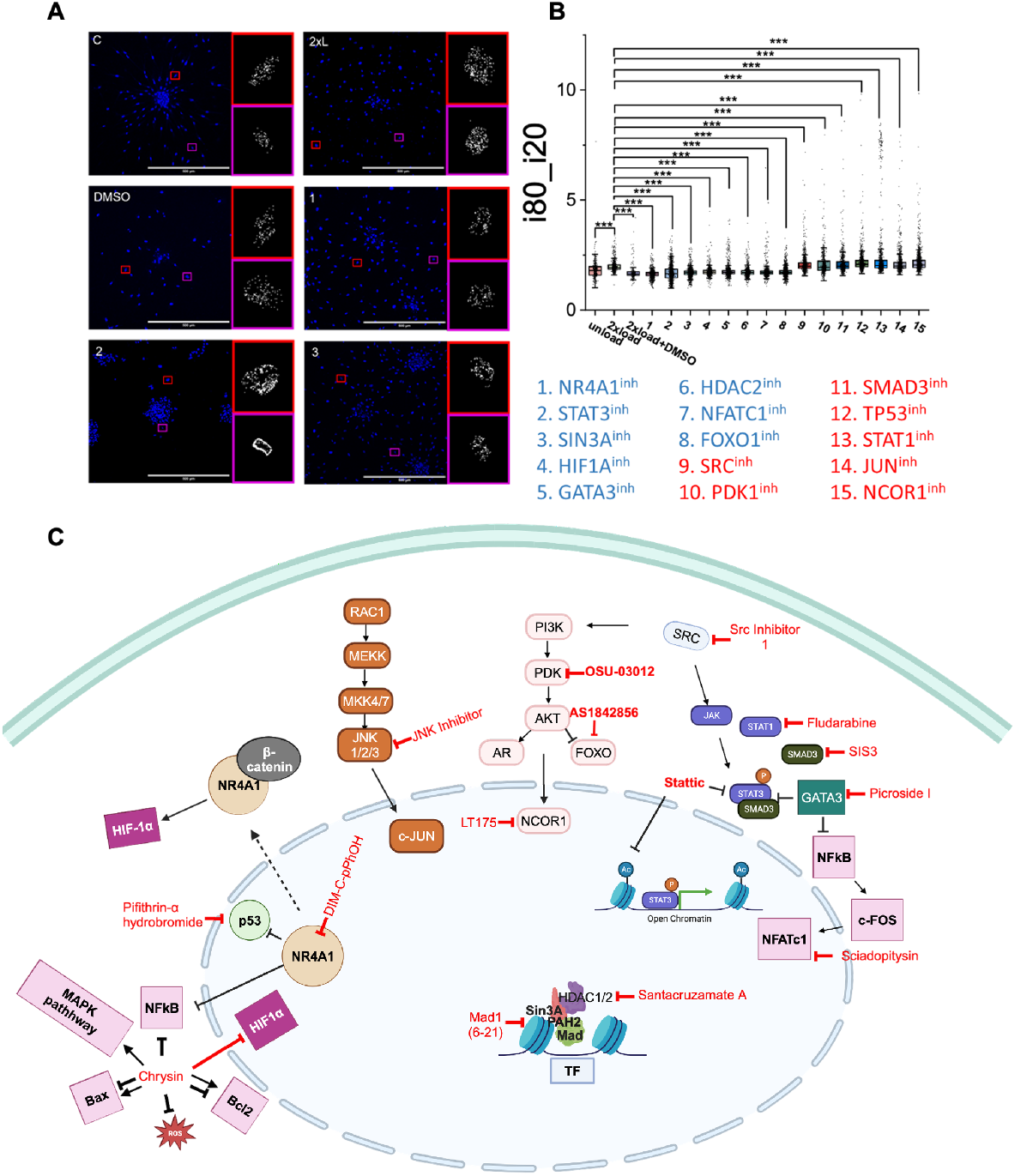
Chromatin condensation changes as a screening readout. (A) Representative DAPI stained image showing heterochromatin distribution in cells from experiments in Figure3. Insert: Ostu thresholded dense chromatin regions were shown. (scale bar 100 μm) (B) i80_i20 represents the ratio of heterochromatin and euchromatin. It is the ratio of the 80%-to-20%-tile of the intensity distribution. P values were calculated by unpaired, two-tailed Student’s t test, compared to the 2xload group. *P<0.05; **<0.01; ***P<0.001; No asterisks means not significant. Blue compounds: low level of i80_i20 compared to 2xload group. Red compounds: higher level than 2xload group. (C) Schematic illustration of inhibitor targets. NR4A1 (Nuclear Receptor Subfamily 4, Group A, Member 1); STAT3 (Signal Transducer and Activator of Transcription 3); SIN3A (SIN3 Transcription Regulator Family Member A); HIF1A (Hypoxia-Inducible Factor 1 Alpha); GATA3 (GATA Binding Protein 3); HDAC2 (Histone Deacetylase 2); NFATC1 (Nuclear Factor of Activated T-cells, Cytoplasmic 1); FOXO1 (Forkhead Box O1); SRC (Proto-oncogene tyrosine-protein kinase Src); PDK1 (Phosphoinositide-dependent kinase-1); SMAD3 (Mothers Against Decapentaplegic Homolog 3); TP53 (Tumor Protein P53); STAT1 (Signal Transducer and Activator of Transcription 1); JUN (c-Jun); NCOR1 (Nuclear Receptor Co-repressor 1).

### The effect of inhibitors on chromatin condensation during force induced fibroblast activation

Apart from migration assay as described before, the chromatin condensation state, which reflects the global gene expression state as described in our lab’s previous works, was also measured as a screenable parameter (72). As heterochromatin is more condensed and euchromatin is more loosely organized, DAPI staining reveals the brighter regions as heterochromatin. To quantify chromatin condensation, we used the Otsu method to identify heterochromatin foci and analyze their distribution, as shown in Figure S4A. In Figure 4A, we observed that in the control group, heterochromatin was largely absent near the nuclear envelope. However, under compressive load, heterochromatin began to accumulate around the nuclear envelope.

Since the inhibitors for STAT3, FOXO1, and PDK1 also suppressed cell migration, we noted the appearance of large heterochromatin dots near the nuclear envelope in these groups, possibly suggesting the formation of SASP (senescence-associated secretory phenotype) loci.

Our previous work demonstrated that the i80 by i20 ratio could be used to estimate the relative abundance of heterochromatin to euchromatin. This ratio is the 80th-to-20th percentile of the intensity distribution. As shown in Figure 4B, cells under the first eight inhibitor treatment exhibited a lower heterochromatin ratio, while cells under the remaining seven inhibitor treatment showed higher heterochromatin levels. Among these seven inhibitors, SRC, SMAD3, TP53, and JUN inhibitors showed increased migration areas compared to the 2x load group (Figure 3A and 3B). These results suggest that inhibiting SRC, SMAD3, TP53, and JUN enhances cellular migration and increases heterochromatin content.

## Discussion

Our previous study demonstrated that compressive force could shift aged cells toward activation or rejuvenation. In this study, through selective TF inhibitor screening, we identified that STAT3, FOXO1, and PDK1 strongly inhibit cell migration. STAT3, part of the JAK-STAT signaling pathway, functions as a signal transducer and activator (29). Several studies have linked STAT3 to mechanical force responsiveness (30, 31). One study found that STAT3 activation is essential for myofibroblast activation (32). Another suggested that constitutive STAT3 activation protects fibroblasts from UV-induced apoptosis (33). However, other studies have associated STAT3 activation with cellular senescence or fibrosis (34–36). This discrepancy may arise from differences in signaling pathways, subcellular localization, epigenetic modifications, feedback loops, and the duration or intensity of STAT3 activation. Specifically, we hypothesize that compressive force induced STAT3 activation is transient and adaptive, whereas biochemical activation is prolonged or excessive. Mechanical stimuli may induce chromatin changes that allow for controlled STAT3 activity, limiting harmful gene expression. Conversely, biochemical activation might trigger widespread gene expression, leading to detrimental outcomes like cellular stress or oncogenic transformation.

Interestingly, one study reported that inhibition of ERK1/2 by PD98059 prevented the serine phosphorylation of STAT3 (37). In our model, both the ERK inhibitor (PD98059) (71) and STAT3 inhibitor (Stattic) led to reduced cell migration, suggesting that the ERK-STAT3 signaling axis may play a role in this process. In addition, since FOXO1 is downstream of PDK1, we focus our analysis on FOXO1. FOXO1, a mechanosensitive transcription factor belonging to the FOXO family, has been implicated in longevity regulation via insulin signaling (38, 39). Overexpression of FOXO1 has been linked to enhanced memory and metabolic fitness (40). One study demonstrated that FOXO1 can be phosphorylated by AKT, leading to its interaction with IQGAP1, which impedes IQGAP1-dependent phosphorylation of ERK1/2 (pERK1/2) (41). In conclusion, based on the current data, we propose that the FOXO1-pERK-STAT3 signaling axis plays a key role in fibroblast activation induced by compressive force.

In the normal state, DNA within the nucleus exerts an outward entropic force due to its coiling, while interactions between DNA, histones, and non-histone proteins generate an inward enthalpic force (10). Under compressive force conditions, the nucleus deforms, resulting in the activation of various biochemical and biophysical signals to alter chromatin condensation. Thus, we use heterochromatin content as another readout for the drug screening. We found that inhibition of SRC, SMAD3, TP53, and JUN enhances cellular migration and increases heterochromatin content under compressive force conditions. SRC kinase activity is known to be critical for focal adhesion turnover and cell motility through the tyrosine phosphorylation of FAK (42). One study has shown that the SMAD complex and JUN can be recruited to the enhancer region of SRC to regulate TGF-β-induced SRC expression (43). Interestingly, in our model, inhibiting SRC, SMAD3, and JUN did not significantly affect cell migration, which aligns with the results from our FAK inhibitor group (71). This suggests that compressive force-induced cell migration in our system may be FAK-independent. Additionally, p53 has been reported to suppress cell motility (44), which is consistent with the inhibitory effect of p53 on migration observed in our study. These findings indicate that the mechanism driving cell migration under compressive force may bypass traditional pathways such as FAK-SRC signaling, suggesting the involvement of alternative mechanotransduction pathways. Furthermore, studies have shown that condensed chromatin can increase nuclear stiffness and enhance its durability in response to external forces, thereby facilitating cell migration in confined environments (45, 46). This concept aligns with our observation of increased heterochromatin content, which may enhance the mechanical resilience of the nucleus during migration under compressive force.

The identified key transcription factors ERK, STAT3, and FOXO1 from this screening assay may have potential cofactors or upstream factors to regulate fibroblast activation and rejuvenation jointly. SP1, SPI1, GATA2, MYC, HDAC1, RARA, and EP300 could be potential candidates as shown in several TF-target databases and TF enrichment tools, including hTFTarget database, TRUST database and iRegulon tool. SP1 (Specificity Protein 1), the first transcription factor isolated from mammalian cells, plays a key role in detecting and eliminating cells with persistent DNA strand breaks (47, 48). It has also shown potential in cellular reprogramming (49). SPI1 (also known as PU.1, Spleen Focus Forming Virus Proviral Integration Oncogene) is highly expressed in matrix-producing fibrotic fibroblasts (50) and plays a role in regulating tissue extracellular matrix (ECM). Given the reduction of ECM content with aging in fibroblasts (51), SPI1’s role is particularly interesting in the context of anti-aging research. GATA2 (GATA Binding Protein 2) is essential for the maintenance and self-renewal of hematopoietic stem cells (52), highlighting its importance in cellular regeneration. MYC (Myelocytomatosis Oncogene) is one of the four Yamanaka factors that reprogram somatic cells into induced pluripotent stem cells (iPSCs) (53), underscoring its critical role in cellular rejuvenation. HDAC1 (Histone deacetylase 1) has been reported to extend lifespan via its inhibition (54). RARA (Retinoic Acid Receptor Alpha), also known as NR1B1, regulates gene expression by interacting with retinoic acid (55). Retinoids, well known for their anti-aging effects in skincare (56), further suggest that RARA could be relevant in cellular aging and rejuvenation. EP300 (E1A Binding Protein p300) is a histone acetyltransferase involved in transcriptional regulation via chromatin remodeling (57). It is also associated with cellular proliferation, differentiation, and senescence (58), making it a promising target in the context of rejuvenation strategies. In summary, these transcription factors are associated with key biological processes related to aging and rejuvenation, including chromatin remodeling, gene expression, and cell cycle regulation. Some, such as MYC and EP300, are directly implicated in reprogramming cells to a more youthful state, while others, like SP1, GATA2, and RARA, help maintain cellular functions that combat age-related decline. Therefore, these transcription factors and chromatin regulators are highly relevant to cellular rejuvenation and present potential therapeutic targets for anti-aging interventions.

Taken together, our results provide a comprehensive understanding of the interplay between transcription factors, epigenetic regulators, and the cytoskeleton in the context of compressive force-induced activation/rejuvenation of aged fibroblasts. These findings offer valuable insights for drug discovery, disease modeling, and regenerative medicine.

## Data and materials availability

All codes used in this paper are available from the corresponding author upon request. All illustration graphs shown in this study were created by Biorender.com. All data are available in the main text or the supplementary materials.

## Acknowledgments

We thank GVS group members for their comments on the manuscript. This work was supported by the Swiss National Science Foundation grant 310030_208046; China Scholarship Council (Grant Number: 202008440471).

## Notes

### Competing Interest Statement

The authors have declared no competing interest.

