## Supplementary File for "Identifying regulators of aged fibroblast activation in 3D tissue models"

**Supplementary material**

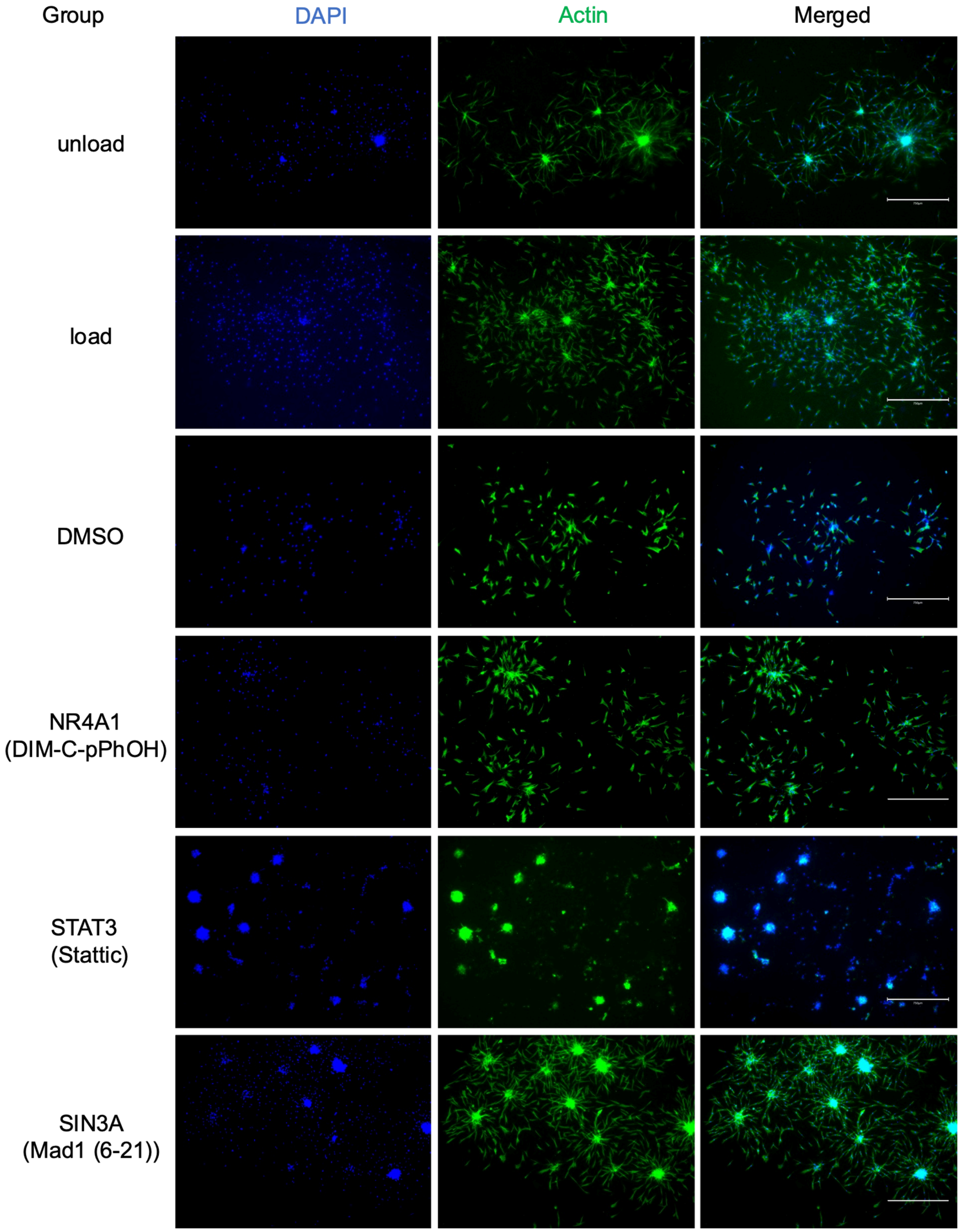

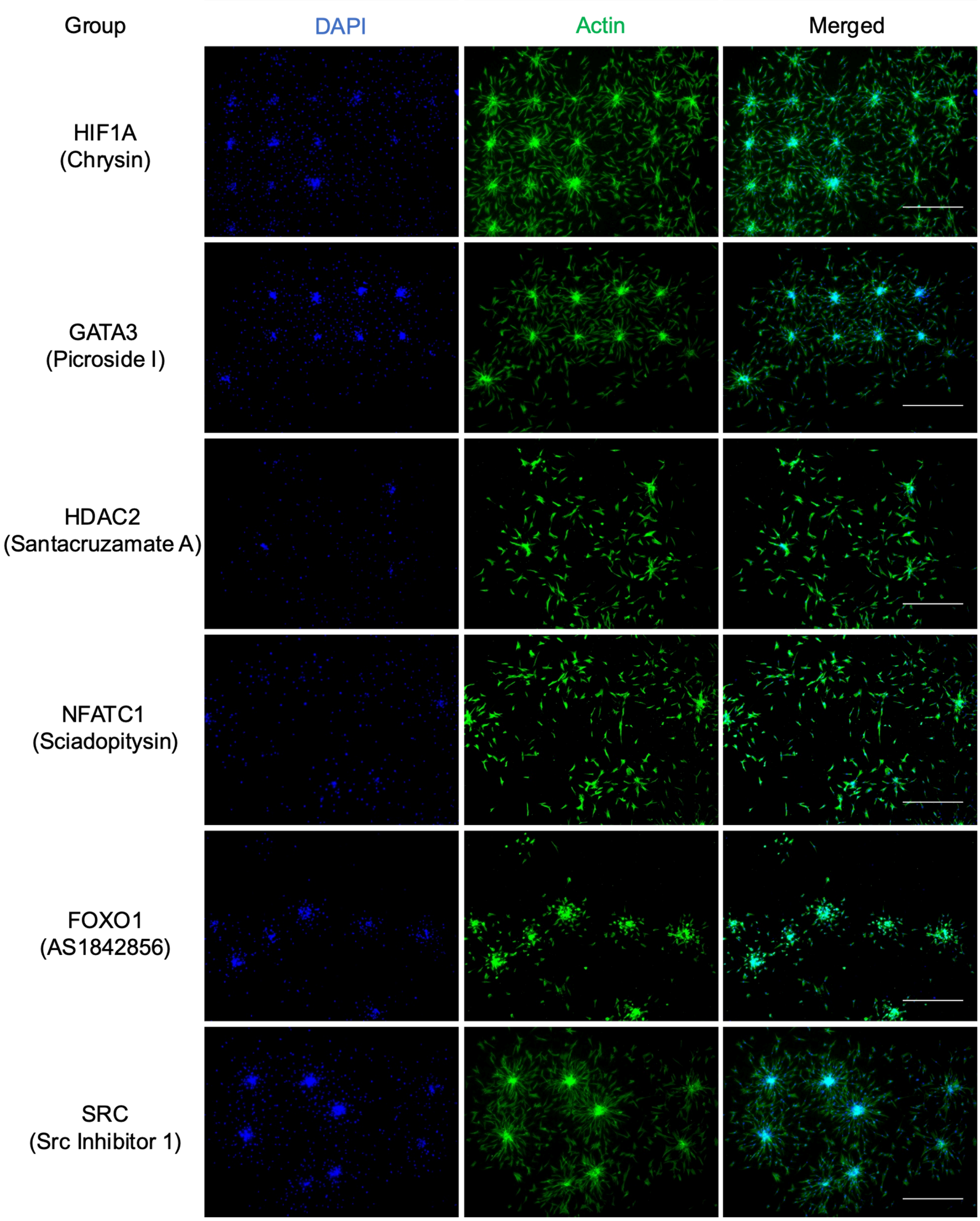

Group
DAPI
Actin
Merged
HIF1A
(Chrysin)
GATA3
(Picroside I)
HDAC2
(Santacruzamate A)
NFATC1
(Sciadopitysin)
FOXO1
(AS1842856)
SRC
(Src Inhibitor 1)

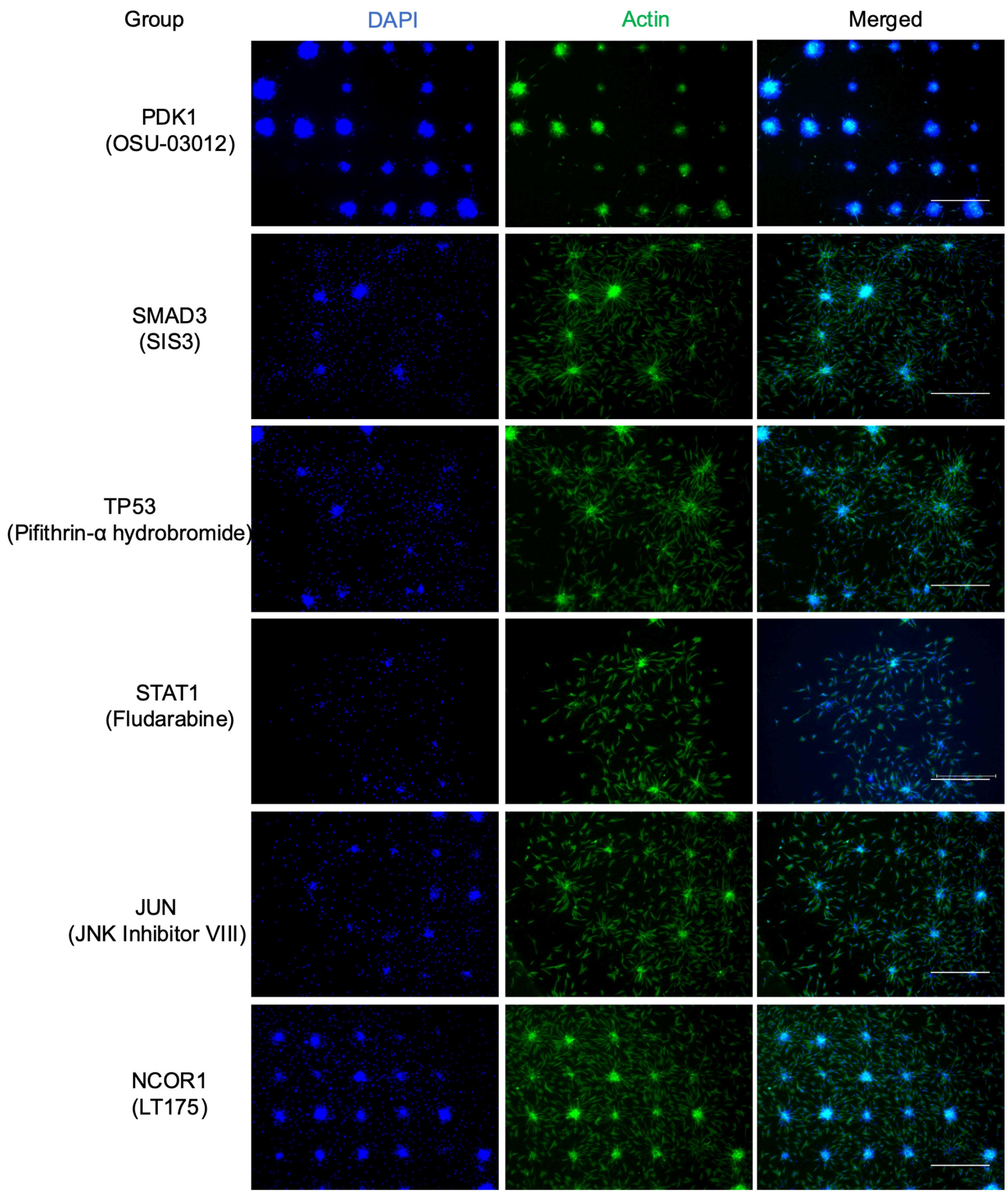


Figure S1. Big areas for TF drug screening (EVOS, 4X objective). (Scale bar, 750 μm). Green color (actin), Blue color (DAPI).

**A**

| TF | Log2FoldChange |
|---|---|
| NR4A1 | 1.24948 |
| CEBPD | 1.39216 |
| KLF9 | 1.07252 |
| ZNF331 | 1.1721 |
| IRF1 | 1.10464 |
| NR6A1 | 1.28187 |

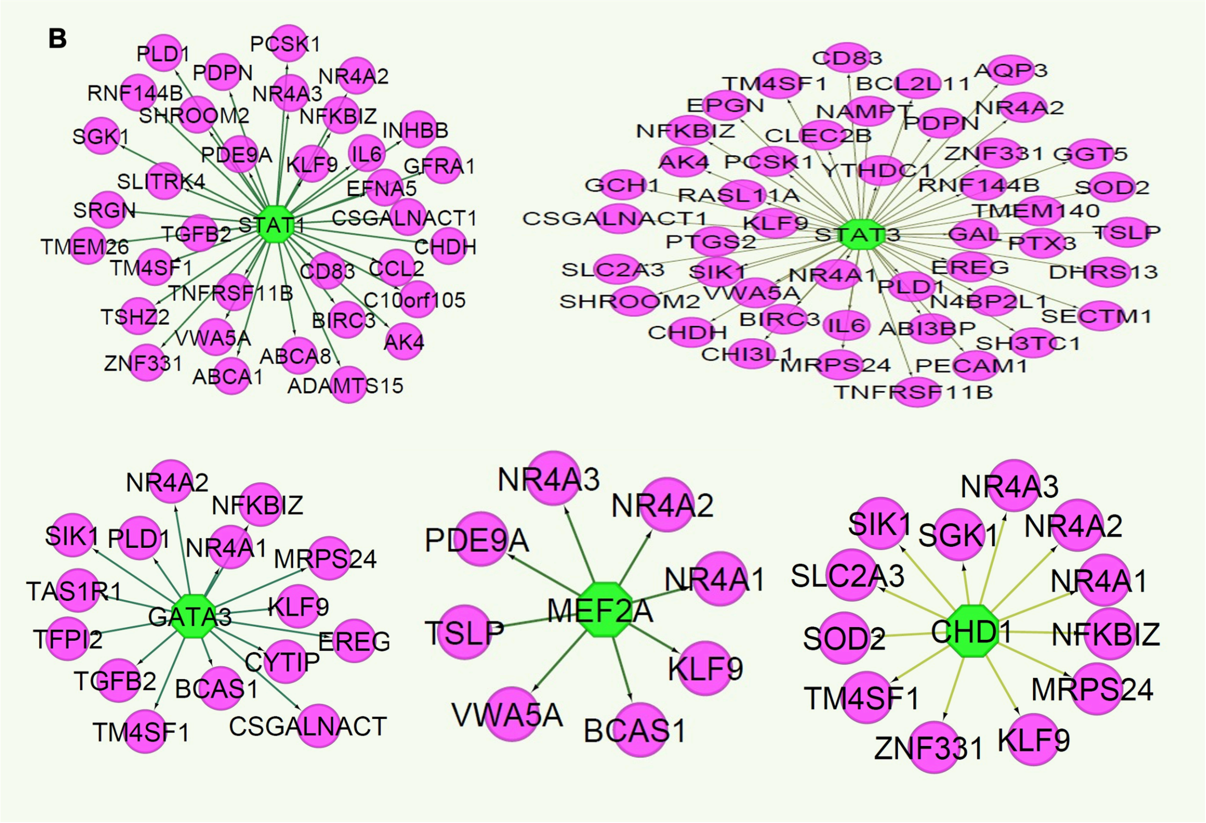


Figure S2. (A) Log2FoldChange of the most differentially expressed transcriptional regulators comparing 2XL and C from the list of pink nodes in Figure 2A. (B) Regulatory network of representative TFs (STAT1, STAT3, GATA3, MEF2A and CHD1) and their target genes, pink nodes represent gene; green nodes represent TF.

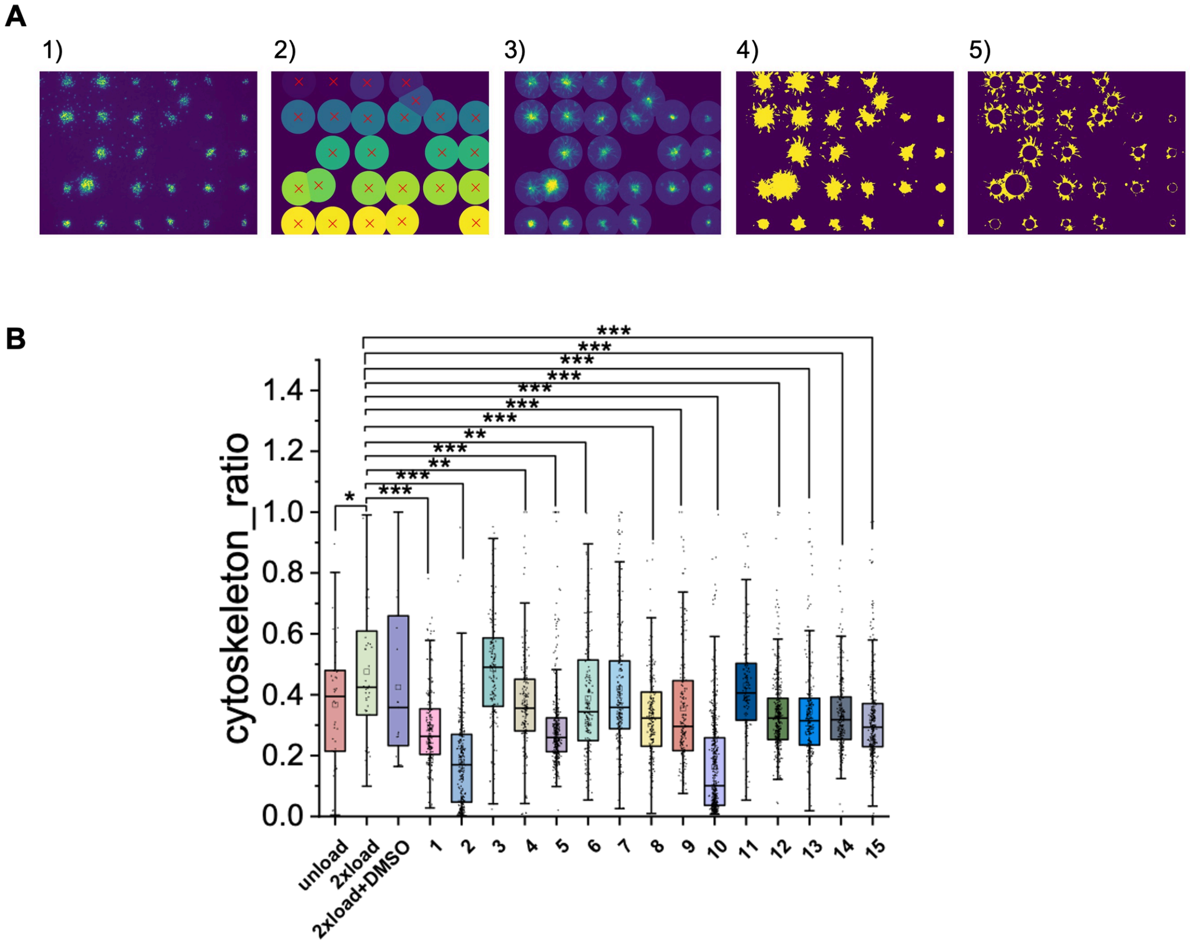


Figure S3. (A) Quantification pipeline for migration assay. DAPI images of spheroids were first segmented to estimate spheroid center. A ROI centered on an inferred spheroid center with diameter of 250 μm was segmented for downstream analysis. Actin stained images within ROI were segmented and the core of the spheroid were estimated using distance_transform_edt from Scipy. (B) Boxplot showing outside of the core of the spheroid but within ROI, the ratio of cytoskeleton area.

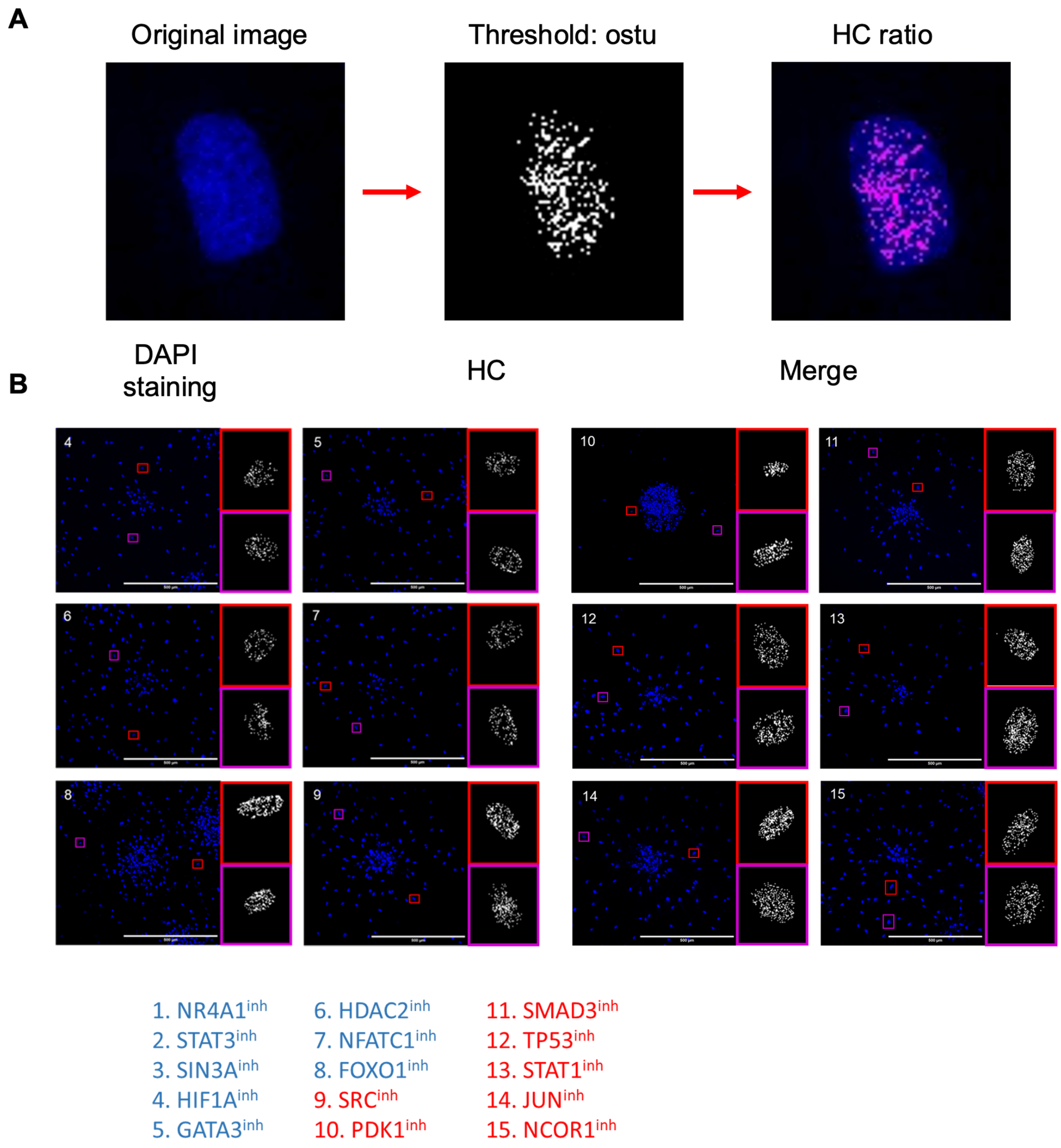


Figure S4. (A) Schematic of measuring the heterochromatin (HC). (B) Representative DAPI stained image showing heterochromatin distribution in cells from experiments in Figure3. Insert: Ostu thresholded dense chromatin regions were shown. Blue compounds: low level of i80_i20 compared to 2xload group. Red compounds: higher level than 2xload group.

**Table S1.** List of inhibitors. (* recommendation from supplier)

| Drug name | Brand & catalogue number | Final concentration |
|---|---|---|
| DIM-C-pPhOH | MCE/HY-112055 | 15 uM (59) |
| Stattic | MCE/HY-13818 | 3.3 uM (60) |
| Mad1 (6-21) | MCE/HY-P3242 | 60nM |
| Chrysin | MCE/HY-14589 | 20 uM (61) |
| Picroside I | MCE/HY-N0407 | 10uM (62) |
| Santacruzamate A | MCE/HY-N0931 | 50uM |
| Sciadopitysin | MCE/HY-N2119 | 10uM (63) |
| AS1842856 | MCE/HY-100596 | 5uM (64) |
| Src Inhibitor 1 | MCE/HY-101053 | 10uM (65) |
| OSU-03012 | MCE/HY-10547 | 20uM (66) |
| SIS3 | MCE/HY-13013 | 10uM (67) |
| Pifithrin-α hydrobromide | MCE/HY-15484 | 10uM (68) |
| Fludarabine | MCE/HY-B0069 | 5uM (69) |
| JNK Inhibitor VIII | MCE/HY-107598 | 50nM (70) |
| LT175 | MCE/HY-121900 | 5uM |
